# Identification of extracellular membrane protein ENPP3 as a major cGAMP hydrolase, cementing cGAMP’s role as an immunotransmitter

**DOI:** 10.1101/2024.01.12.575449

**Authors:** Rachel Mardjuki, Songnan Wang, Jacqueline A. Carozza, Gita C. Abhiraman, Xuchao Lyu, Lingyin Li

**Affiliations:** Department of Biochemistry, Stanford University, Stanford, CA 94305, USA; ChEM-H Institute, Stanford University, Stanford, CA 94305, USA; Arc Institute, Palo Alto, CA 94304, USA; Department of Chemistry, Stanford University, Stanford, CA 94305 USA; Department of Molecular and Cellular Physiology, Stanford University, Stanford, CA 94305, USA; Department of Pathology, Stanford University, Stanford, CA 94305, USA

## Abstract

cGAMP is a second messenger that is synthesized in the cytosol upon detection of cytosolic dsDNA and passed between cells to facilitate downstream immune signaling. ENPP1, an extracellular enzyme, was the only metazoan cGAMP hydrolase known to regulate cGAMP levels to dampen anti-cancer immunity. Here, we uncover ENPP3 as the second and only other metazoan cGAMP hydrolase under homeostatic conditions. ENPP3 has a tissue expression pattern distinct from that of ENPP1 and accounts for all remaining cGAMP hydrolysis activity in mice lacking ENPP1. Importantly, we also show that as with ENPP1, selectively abolishing ENPP3’s cGAMP hydrolase activity results in diminished cancer growth and metastasis of certain tumor types. Both ENPP1 and ENPP3 are extracellular enzymes, suggesting the dominant role that extracellular cGAMP must play as a mediator of cell-cell innate immune communication. Our work clearly shows that ENPP1 and ENPP3 non-redundantly dampen extracellular cGAMP-STING signaling, pointing to ENPP3 as a new target for cancer immunotherapy.

## Introduction

The presence of cytosolic double-stranded DNA (dsDNA) or DNA/RNA hybrids can be a sign of viral infection, cancer, aging, autoimmunity, or tissue damage. 2’3’-Cyclic GMP-AMP (cGAMP) is an important innate immune second messenger that is synthesized in the cytosol by the enzyme Cyclic GMP-AMP Synthase (cGAS) upon detection of these molecular patterns ^1–5^. cGAMP potently activates its intracellular receptor, Stimulator of Interferon Genes (STING), leading to the production of type-I interferons and other interferon-induced genes to mount an innate immune response to the detected threat ^2,6,7^. In addition to signaling within the producing cell, cGAMP is also known to be exported from some producer cells, such as cancer cells, and activate the STING pathway in other cell types within the nearby vicinity ^8–12^. Activation of STING in responder cells, including fibroblasts, macrophages, and endothelial cells, elicit downstream anti-cancer ^8,11,12^ and anti-viral immune responses, but also can exacerbate systemic inflammation and autoimmunity ^13^. Interestingly, while STING activation in cancer cells can promote metastasis ^14,15^, STING activation in bystander cells of the tumor microenvironment delays primary tumor growth and metastasis and is responsible for the curative effect of radiation treatment ^8,12,16–21^. These nuances emphasize the importance of cell type- and tissue-specific regulation of cGAMP levels both inside and outside the cell.

Ectonucleotide pyrophosphatase phosphodiesterase I (ENPP1) was reported 10 years ago as the first metazoan cGAMP hydrolase. ENPP1 is a type-II transmembrane protein with its catalytic domain sitting on the outer surface of the cell membrane and has been shown to regulate extracellular cGAMP levels in the context of cancer, infection, local and systemic inflammation ^8,12,13,22–24^. Intriguingly, other second messengers like cGMP and cAMP can be degraded by 11 different phosphodiesterase families (PDE1 – PDE11), contributing to their cell-, tissue-, and context-specific regulation ^25^. However, it remains unclear whether there are other enzymes capable of degrading cGAMP that may act as a context- or tissue-specific regulator of STING activation.

Here, we set out to address this question by interrogating and isolating any remaining cGAMP hydrolase activity in mice lacking *Enpp1*. While some tissues in *Enpp1* knockout (*Enpp1*^-/-^) mice no longer have detectable cGAMP hydrolase activity, other tissues like the kidney display significant remaining cGAMP hydrolase activity. We identified the responsible enzyme to be ENPP3, another extracellular enzyme and paralog of ENPP1. Mice harboring a point mutant of ENPP3 that specifically abolishes its cGAMP hydrolase activity are more resistant to both primary tumor growth and metastasis in certain cancers. This study offers a new target for cancer immunotherapy as well as new mouse strains that can help elucidate extracellular cGAMP’s roles in health and disease. We detected no remaining activity in mice after disrupting the cGAMP hydrolase activity of both ENPP1 and ENPP3. We conclude that these two enzymes are likely the only two dominant mouse cGAMP hydrolases under homeostatic conditions, suggesting that all degradative regulation of cGAMP occurs extracellularly and that the predominant mode of signaling of cGAMP is through paracrine STING activation.

## Results

### Identification and validation of ENPP3 as a cGAMP hydrolase

*ENPP1* expression levels in cancer patients have been shown to correlate with treatment response and prognosis^12^, and *Enpp1*^-/-^ mice are routinely used to assess the contribution of cGAMP hydrolysis to regulating innate immune activity. However, it remains unclear whether ENPP1 is the sole hydrolase of cGAMP, or whether other enzymes also contribute. To answer this question, we harvested and homogenized various organs from WT or *Enpp1*^-/-^ mice to measure their cGAMP hydrolase activity (**Figure 1A, B**). As expected, tissues where ENPP1 was initially purified from and/or characterized in, including the liver, spleen, and plasma^26^, had negligible cGAMP hydrolase activity remaining after *Enpp1* knockout (**Figure 1A, B**). In addition, ENPP1 also accounts for all the cGAMP hydrolase activity in the brain (**Figure 1B**). However, most *Enpp1*^-/-^ organs exhibited considerable amount of residual cGAMP hydrolase activity, including the kidney, ovary, uterus, cervix, mammary fat pad, lung, and heart. Notably, the kidney has both the most total activity and the highest overall cGAMP hydrolase activity to protein ratio, of which more than half is contributed by the unknown hydrolase(s) (**Figure 1A, B**).

**Figure 1:**
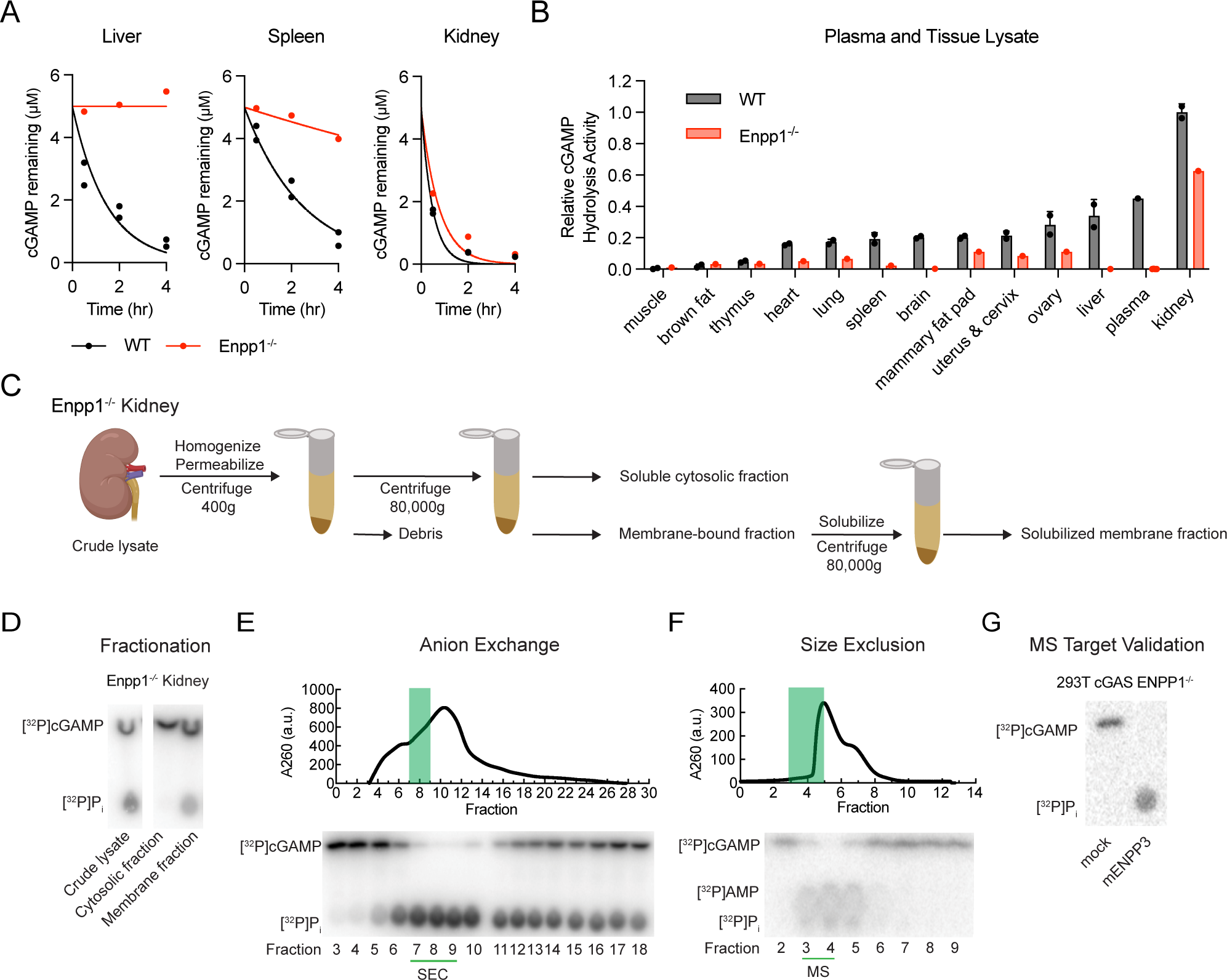
Identification and validation of ENPP3 as a cGAMP hydrolase (A) Example kinetic analysis of cGAMP degradation doped with ^32^P-cGAMP (pH 7.5, 37°C) by organ lysate harvested from WT and *Enpp1*^-/-^ mice. Starting material and product was measured by thin-layer chromatography (TLC) and visualized by autoradiography and quantified using ImageJ. (B) Rate constant of cGAMP degradation in all organs determined as in example kinetic analysis*. n* = 2 animals for WT, *n* = 1 animals for *Enpp1*^-/-^, mean ± SD with some error bars too small to visualize. (C) Schematic for fractionation of ENPP3 from kidneys from *Enpp1*^-/-^ mice. (D) cGAMP degradation activity of fractions indicated in Figure 1C at physiological concentrations of divalent ions at Tris pH 7.5, 37°C. Chromatography trace and cGAMP degradation activity from (E) HiTrap Q HP anion exchange fractionation and (F) size exclusion Superose 12 fractionation of *Enpp1*^-/-^ mouse kidney lysate. Green bar highlights fractions with highest cGAMP hydrolysis activity/total protein concentration ratio. (G) cGAMP degradation by 293T *ENPP1^-/-^* cells transfected with mENPP3 (pH 7.5, 37°C).

Therefore, we focused on *Enpp1*^-/-^ kidney to isolate the unknown hydrolase(s) using serial fractionation (**Figure 1C**). We measured cGAMP hydrolase activity in the crude lysate, cytosolic fraction, and membrane fraction of *Enpp1*^-/-^ kidney lysates (**Figure 1C, D**). Only the membrane fraction, not the cytosolic fraction, had detectable levels of cGAMP hydrolase activity, suggesting the lack of intracellular cGAMP hydrolase activity in kidneys in the condition tested (**Figure 1D**). To further isolate the hydrolase(s), we subjected the solubilized membrane fraction to sequential anion exchange and size-exclusion chromatography (SEC) (**Figure 1E, F**). We analyzed the composition of size-exclusion fraction 4 with the highest cGAMP hydrolase activity to protein ratio using mass spectrometry (MS). Size-exclusion fraction 3, which has less cGAMP hydrolase activity than fraction 4, was also submitted for MS analysis.

Among the top hits, ENPP3, a paralog of ENPP1, was the fourth most abundant protein detected in fraction 4 (**Supplemental Table 1**). ENPP3 was less abundant in fraction 3 as expected, further indication of ENPP3 as the unknown cGAMP hydrolase in *Enpp1^-/-^* kidney. Among the seven members of the ENPP family (ENPP1 – ENPP7), ENPP1 and ENPP3 are the most closely related both evolutionarily and structurally (**Figure S1A, B**) ^27^. They are encoded only 60 kb apart on chromosome 6 and 10 of humans and mice, respectively. Like ENPP1, ENPP3 is a single-pass transmembrane protein with its catalytic domain located on the extracellular side of the plasma membrane. To validate that ENPP3 is indeed a cGAMP hydrolase, we transiently transfected mouse ENPP3 (mENPP3) expressing construct into 293T cGAS *ENPP1*^-/-^ cells ^8^. The mENPP3 expressing cell lysate degraded cGAMP efficiently, whereas the untransfected control did not (**Figure 1G**). Together, these findings point to ENPP3 as the second mammalian cGAMP hydrolase.

### ENPP3 is an efficient hydrolase of cGAMP at physiological pH

Next, we sought to biochemically characterize ENPP3’s cGAMP hydrolase activity. ENPP1 degrades cGAMP extracellularly, as overexpressing ENPP1 in cell lines diminished extracellular cGAMP without affecting intracellular cGAMP levels ^8^. ENPP3, in contrast, has previously been shown to be active intracellularly in the lumen of the Golgi apparatus. In the Golgi, ENPP3 hydrolyzes UDP-GlcNAc to inhibit the function of acetylglucosaminyltransferase GnT-IX and thereby affects global cell surface protein glycosylation ^28^.

Because cGAMP is synthesized in the cytosol and has not been shown to be transported into the Golgi, we hypothesize that only the plasma membrane and therefore extracellular pool of ENPP3 degrades cGAMP. To test this hypothesis, we transiently transfected mENPP3 and human ENPP3 (hENPP3) expressing constructs into 293T cGAS *ENPP1*^-/-^ cells. Like ENPP1, overexpression of mENPP3 and hENPP3 decreased extracellular cGAMP without affecting intracellular cGAMP, suggesting only extracellular ENPP3 is a hydrolase for cGAMP (**Figure 2A**).

**Figure 2:**
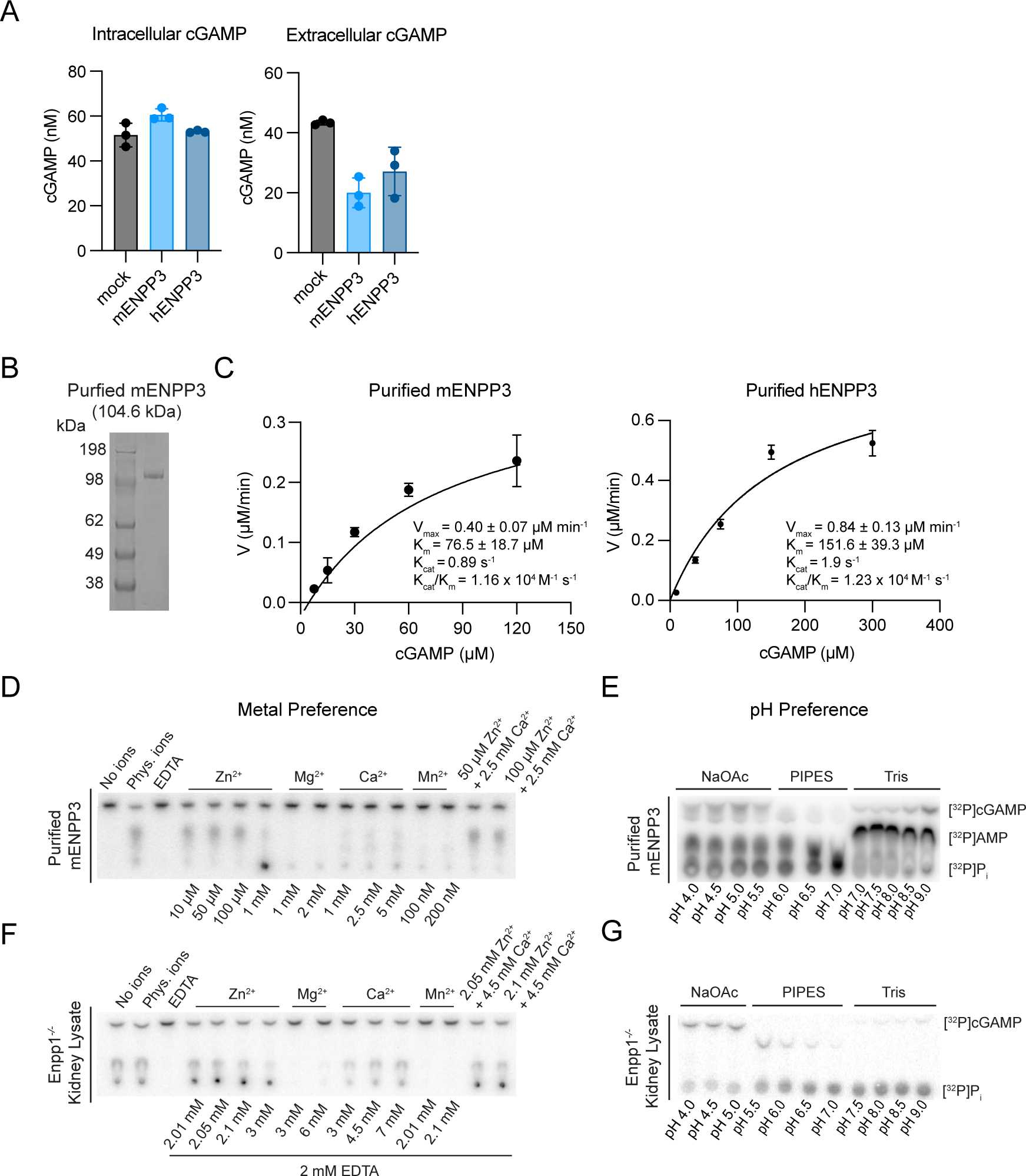
ENPP3 is an efficient hydrolase of cGAMP at physiological pH (A) Intracellular and extracellular cGAMP quantified by cGAMP ELISA in ENPP3-transiently transfected 293T *ENPP1^-/-^*cells. (B) Purified recombinant mouse ENPP3 (mENPP3) visualized by Coomassie gel. (C) Kinetics of cGAMP hydrolysis of purified mouse and human ENPP3. Mean ± SEM (*n* = 3 independent experiments). (D, F) Divalent ion preference (measured in Tris pH 7.5) and (E, G) pH preference (measured in physiological concentrations of divalent ions) of *Enpp1*^-/-^ mouse kidney and purified mENPP3.

Using purified recombinant mENPP3 (**Figure 2B**), we determined that ENPP3 efficiently hydrolyzes cGAMP (*K_m_* = 76.5 ± 18.7 μM, *k*_cat_ = 0.89 s^-1^) at pH 9.0 (**Figure 2C**). hENPP3 also hydrolyzes cGAMP efficiently (*K*_m_ = 151.6 ± 39.3 μM, *k*_cat_ of 1.9 s^-1^, and *k*_cat_ /*K*_m_ = 1.23 x 10^4^ M^-1^s^-1^) (**Figure 2C**), albeit approximately 10-fold less efficient than hENPP1 (*K*_m_ = 15 μM, *k*_cat_ = 4 s^-1^, and *k*_cat_ /*K*_m_ = 2.75 x 10^4^ M^-1^s^-1^) in the same condition at pH 9.0 ^26^. We then determined the optimal ion and pH conditions for degradation of cGAMP by ENPP3. It is known ENPP3 requires two Zn^2+^ ions for catalysis and Ca^2+^ ions for structural integrity^29,30^. We confirmed this ion preference in recombinant mENPP3, where addition of Zn^2+^ and Ca^2+^ but not the other divalent ions (Mg^2+^ and Mn^2+^) increased mENPP3’s hydrolase activity (**Figure 2D**). ENPP1 peaks in activity at pH 9.0 ^26^; however, ENPP3 prefers neutral and alkaline pHs, with a peak in activity around pH 7.0 (**Figure 2E**). Strikingly, we also found the same divalent cation and pH preference of 7.0 in the EDTA-treated *Enpp1*^-/-^ mouse kidney lysate, strongly suggesting that ENPP3 accounts for most if not all the remaining cGAMP hydrolase activity in mice (**Figure 2F, G**).

### ENPP3 is the only other cGAMP hydrolase in mice

To determine ENPP3’s contribution to cGAMP hydrolase activity in tissues and to investigate ENPP3’s roles *in vivo*, we next sought to generate *Enpp3* knockout (*Enpp3*^-/-^) mice. However, like ENPP1 ^31^, ENPP3 not only hydrolyzes extracellular cGAMP, but also extracellular ATP. In addition, ENPP3 also hydrolyzes UDP-GlcNAc ^28^. While ENPP1 degrades ATP to maintain calcium homeostasis and prevent arterial and joint calcification, ENPP1 degrades cGAMP to prevent systemic inflammation, but inadvertently promotes cancer and viral infections ^12,13^. We thus created mice harboring the H362A ENPP1 point mutation (*Enpp1^H362A^*) ^13^. The H362A mutation abolishes ENPP1’s activity towards cGAMP but not ATP, allowing us to study the divergent roles of ENPP1 in regulating these two substrates ^12,13^. These findings demonstrate the need to isolate ENPP3’s roles in regulating cGAMP versus other substrates.

As ENPP1 and ENPP3 have similar catalytic sites, we identified H329 as the histidine in ENPP3 analogous to H362 in ENPP1. *In vitro* studies confirmed that the H329A mutation in mENPP3 completely abolishes its hydrolase activity towards cGAMP while preserving its activity towards ATP (**Figure 3A, B**). We confirmed that the H329A mENPP3 mutation also retains its hydrolase activity towards UDP-GlcNAc (**Figure S2**).

**Figure 3:**
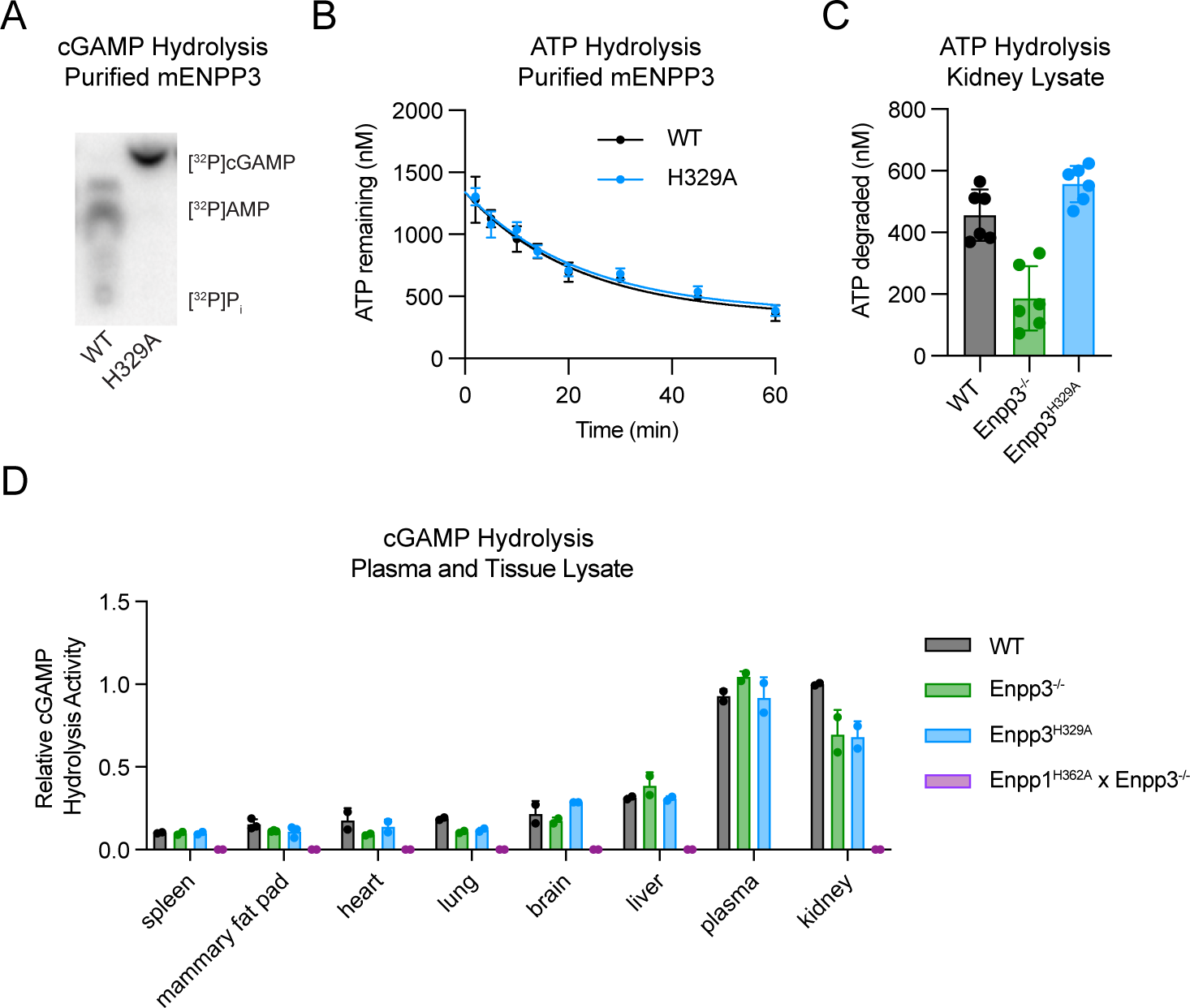
ENPP3 is the only other hydrolase in mice (A) cGAMP degradation by purified WT and mENPP3 H329A. (B) Kinetics of ATP degradation by purified WT and mENPP3 H329A measured by CellTiter-Glo assay. Mean ± SEM (*n* = 3). (C) ATP degradation by kidney lysates from mice of indicated genotypes quantified by CellTiter-Glo assay. Mean ± SEM (*n* = 3 animals and n=2 independent experiments for each animal). (D) Rate constant of cGAMP degradation in organ lysate determined as in Figure 1(A). (n=2 animals per genotype; n=3 animals for mammary fatpad only).

Using CRISPR-based homologous recombination, we generated *Enpp3*^-/-^ and homozygous *Enpp3*^H*329*A^ mice on the C57BL/6 genetic background (**Figure S3A-E**). We confirmed that kidney lysate from *Enpp3*^H*329*A^ mice degraded ATP similarly to that of WT mice, indicating that ENPP3 is still capable of degrading ATP with the H329A mutation. However, *Enpp3^-/-^* kidney lysate degraded less than half the ATP of the WT mice, indicating that ENPP3 ATP hydrolysis activity was abolished as expected in the *Enpp3*^-/-^ mice (**Figure 3C**). In tissues with considerable cGAMP hydrolase activity remaining after *Enpp1* knockout, such as the kidney, lung, heart, and mammary fat pad (**Figure 1B**), we observed decreased cGAMP hydrolase activity from both *Enpp3*^-/-^ and *Enpp3*^H*329*A^ mice compared with WT (**Figure 3D**). In contrast, in the plasma, liver, brain, and spleen where cGAMP hydrolase activity was primarily attributable to ENPP1 (**Figure 1B**), *Enpp3*^-/-^ and *Enpp3*^H*329*A^ did not impact cGAMP hydrolase activity (**Figure 3D**). Together, we found that using H329A, the analogous separation-of-function mutation to ENPP1’s H362A, we were able to selectively abolish ENPP3 cGAMP hydrolase activity *in vitro* and *in vivo*.

To definitively determine if ENPP1 and ENPP3 are the only cGAMP hydrolases in mice across tissues, we generated mice lacking both ENPP1 and ENPP3 cGAMP hydrolysis activity. *Enpp1* and *Enpp3* are linked genes due to their close proximity to each other, so we could not cross our *Enpp1*^H362A^ mice to our *Enpp3^H329A^* mice to generate *Enpp1*^H362A^ x *Enpp3*^H329A^ mice. We thus performed a second round of CRISPR-based editing of *Enpp3* on the *Enpp1*^H362A^ background. The rate of successful homology-directed repair in mouse embryos has been reported to be as low as ∼1-2%^32^, so with only ten founder mice we were unable to obtain any *Enpp1*^H362A^ x *Enpp3*^H329A^ mice and instead generated homozygous *Enpp1*^H362A^ x *Enpp3*^-/-^ mice.

Interestingly, *Enpp1*^H362A^ x *Enpp3*^-/-^ mice do not exhibit joint calcification phenotype as *Enpp1^-/-^* mice, suggesting that ENPP1’s ATP hydrolysis activity alone is responsible for preventing joint calcification in mice. Tissues harvested from *Enpp1*^H362A^ x *Enpp3*^-/-^ mice displayed no detectable residual cGAMP hydrolase activity, confirming that ENPP3 is the only other dominant hydrolase in mice aside from its paralog ENPP1 (**Figure 3D, S4**).

### ENPP1 and ENPP3 have different expression patterns that influence their effect on cancer

While ENPP1 and ENPP3 are related enzymes with similar biochemical functions, we probed whether they have distinct tissue expression patterns. Indeed, in data plotted from the Human Protein Atlas, we found that while *ENPP1* and *ENPP3* are expressed to similar levels in tissues such as breast and prostate, their expression levels are orthogonal in many other tissues. *ENPP1* and *ENPP3* expression levels are orthogonal in many tissues. For example, placenta and liver predominantly express *ENPP1* while small intestine and basophils predominantly express *ENPP3* (**Figure 4A**)^27^. Furthermore, ENPP1 accounts for virtually all the cGAMP hydrolase activity in mouse plasma: *Enpp1*^-/-^ mouse plasma no longer have any detectable cGAMP hydrolase activity (**Figure 1B, 3D**).

**Figure 4:**
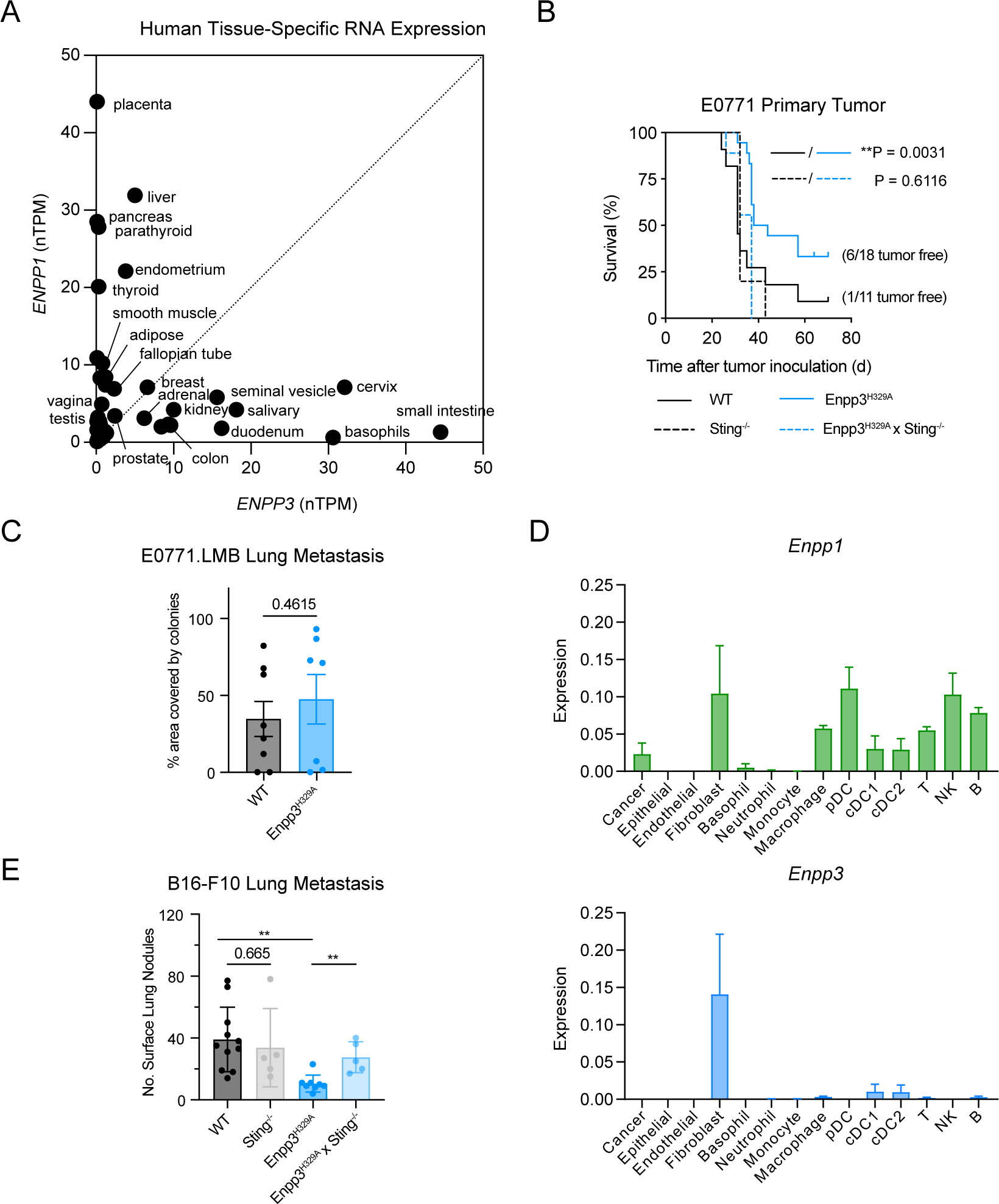
*Enpp3*^H362A^ mice are resistant to melanoma lung metastasis (A) Expression of ENPP1 and ENPP3 from single cell RNA sequencing data from human endometrium tissue. Data from the Human Protein Atlas (https://www.proteinatlas.org/). (B) E0771 cells were orthotopically injected into the indicated genetic background on day 0: WT (n=11), *Sting^-/-^* (n=5), *Enpp3^H329A^* (n=18), and *Enpp3^H329A^Sting^-/-^* (n=9). Mice were euthanized when tumor volume reached 1000 mm^3^. P-values for Kaplan-Meier curves were determined by log-rank Mantel-Cox test. (C) E0771.LMB cells were intravenously injected into the tail vein of mice with the indicated genotype: WT (n=8), *Enpp3^H329A^* (n=7)) on day 0. On day 30, lungs were harvested from mice, homogenized and plated in the presence of puromycin to select for metastatic E0771 cells. Metastasis of each mouse was scored following staining of the E0771 cells by bromophenol blue and an unpaired t-test was used to calculate significance. (D) Bar graphs of expression of *Enpp1* and *Enpp3* in indicated cell types from single-cell RNA sequencing of 4T1 mouse breast cancer pulmonary metastasis. (E) B16 melanoma cells were intravenously injected into the tail vein of mice with the indicated genotype on day 0: WT (n=11), *Sting^-/-^* (n=5), *Enpp3^H329A^* (n=8), *Enpp3^H329A^Sting^-/-^* (n=5). On day 20, lungs were harvested from mice and fixed. Surface metastases were counted and graphed using an unpaired t-test to calculate significance.

In disease settings, *ENPP1* is known to be highly expressed in many human cancer types including breast cancer and glioblastoma^33,34^. We and others previously showed that ENPP1 degrades cancer-produced extracellular cGAMP to dampen downstream STING signaling in stromal cells and tumor-infiltrating immune cells ^8,12,23^. Here, we asked whether *ENPP3* is also highly expressed in some cancer types. We analyzed patient data from the Cancer Genome Atlas (TCGA) to compare *ENPP1* and *ENPP3* expression across cancer types (**Figure S5**). Similar to the distinct expression patterns of *ENPP1* and *ENPP3* in normal human tissues, they are also differentially expressed in human cancers. In some cancer types such as advanced thyroid carcinoma (THCA), only *ENPP1* is expressed; in some cancer types such as colon adenocarcinoma (COAD), kidney renal clear cell carcinoma (KIRC), prostate adenocarcinoma (PRAD), and uterine corpus endometrial carcinoma (UCEC), *ENPP3* is uniquely expressed; and in many breast invasive carcinoma (BRCA) patients, both *ENPP1* and *ENPP3* are expressed.

Because of its expression in normal and cancerous breast tissue, we next investigated if ENPP3, like ENPP1, is an innate immune checkpoint that promotes breast cancer by dampening the cGAMP-STING pathway ^12^. Indeed, in the orthotopic E0771 breast cancer model, mice harboring the cGAMP hydrolysis-deficient *Enpp3* gene (*Enpp3*^H329A^) were significantly more resistant to tumor initiation and growth than WT mice: 6/18 mice remained tumor-free by the end of the 10-week study, compared to 1/11 tumor-free WT mice (**Figure 4B**). This effect was entirely STING-dependent, as *Enpp3*^H329A^ x *Sting*^-/-^ mice have similar tumor progression as WT mice and *Sting*^-/-^ mice. These results mirror our previous observation that mice harboring the cGAMP hydrolysis-deficient *Enpp1* gene (*Enpp1*^H362A^) exhibit STING-dependent delayed primary tumor growth in the same orthotopic E0771 breast cancer model ^12^.

We previously demonstrated that the expression of *Enpp1* is deterministic of breast cancer metastasis into the lungs and other organs ^12^. We thus hypothesized that ENPP3 would not affect breast cancer lung metastasis. However, since both *Enpp1* and *Enpp3* are expressed in mouse lungs, we formally investigated the role ENPP3 plays in breast cancer lung metastasis. In this experiment, E0771.LMB breast cancer cells are introduced into circulation through tail vein injections to induce pulmonary metastasis. As expected, unlike what we observed with *Enpp1*^H362A^ mice, we did not observe appreciable differences in lung metastatic burden between *Enpp3*^H329A^ and WT mice 30 days after cancer cell injection (**Figure 4C**).

To understand the immunological mechanism of the differential effect of ENPP1 and ENPP3 in breast cancer lung metastasis, we mined our previous single-cell RNA sequencing dataset of 4T1 mouse breast cancer pulmonary metastasis to see how *Enpp1* and *Enpp3* are differentially expressed in the tumor microenvironment. In this dataset, *Enpp1* is highly expressed on infiltrating immune cells such as macrophages, pDCs and natural killer cells, as well as tumor-associated fibroblasts (**Figure 4D**)^12^. However, *Enpp3* is not expressed on these infiltrating immune cells, but rather on lung fibroblasts. Breast cancers are known to be heavily infiltrated by immune cells, perhaps explaining the major role of ENPP1 in breast cancer metastasis ^35^.

Given fibroblasts’ general role in supporting tumor growth ^36^, we hypothesize that ENPP3 might play a bigger role in cancer types with less infiltration of immune cells. To test this hypothesis, we chose the B16-F10 melanoma model which is known to have fibroblast involvement but very little immune infiltration ^37^. B16-F10 melanoma cells were introduced into circulation through tail vein injections to induce pulmonary metastasis. Indeed, thirty days after inoculation, we found that *Enpp3*^H329A^ mice have significantly fewer surface lung nodules than WT mice (**Figure 4E**). The effect is STING-dependent, as *Enpp3*^H329A^ x *Sting*^-/-^ mice have similar number of lung nodules as WT and *Sting*^-/-^ mice. These data suggest that in certain cancer types ENPP3, like ENPP1, is an innate immune checkpoint that promotes metastasis by degrading cGAMP and dampening STING pathway activation.

## Discussion

cGAMP is the most recently discovered mammalian second messenger, and despite growing evidence of its cornerstone role in cancer immunology, anti-viral responses, and autoimmunity, our understanding of its physiology in complex tissues and systems remains incomplete. Other second messengers like cAMP and cGMP have eleven different phosphodiesterases (PDEs), making ENPP1 unlikely to be the sole metazoan hydrolase of cGAMP. Here we report the discovery of ENPP3 as a cGAMP hydrolase. Our *Enpp1*^H362A^ x *Enpp3*^-/-^ mice have no detectable cGAMP hydrolase activity, indicating that ENPP1 and ENPP3 are the dominant hydrolases of cGAMP.

We cannot rule out the possibility that other cGAMP hydrolases, intracellular or extracellular, could be induced or activated upon detection of a pathogen or other stimulus. Indeed, while this paper was under review, the discovery of SMPDL3A, a third cGAMP hydrolase, was reported^38^. SMPDL3A is a secreted enzyme whose expression is induced by liver X receptor ligands. This induction is required for the activity of SMPDL3A, as we could not detect any evidence of its activity despite thorough investigation of our *Enpp1*^H362A^ x *Enpp3*^-/-^ mice (**Figure 3D, S4**). As such, the physiological relevance of degradation of cGAMP by SMPDL3A, particularly in the cancer setting, remains unknown. Interestingly, like ENPP1 and ENPP3, SMPDL3A is an extracellular enzyme.

Other second messengers such as cAMP and cGMP are only reported to have intracellular functions and intracellular regulation. In contrast, cGAMP has only been found to have extracellular regulation with a large number of known cGAMP transporters ^9,11,21,39–42^ and hydrolases that are all extracellular. It thus seems reasonable that the grand majority of cGAMP regulation would occur in the extracellular space. Indeed, we have shown that the presence of cytosolic DNA leads to the production of extracellular cGAMP to mediate immunity in a STING-dependent manner ^8,11,13^. Together, we propose a model in which cGAMP’s main physiological function is that of a paracrine immunotransmitter ^8,13,43,44^, and the differential expression of *Enpp1* and *Enpp3* regulate how much cGAMP cells emit and receive.

Our studies of primary breast tumor growth as well as melanoma lung metastasis suggest that like ENPP1, ENPP3 is an innate immune checkpoint that should be targeted to boost anti-cancer immunity. ENPP1 and ENPP3 are similar but distinct enzymes, both with regards to their expression patterns as well as their biochemical properties. Similar distinct expression patterns have been exploited in PDE inhibition, in which drugs targeting different PDEs have different indications depending on their tissue expression. PDE4 is predominantly expressed in the lungs, and its inhibitor Roflumilast treats severe chronic obstructive pulmonary disease (COPD); PDE5 is predominantly expressed in vascular and trabecular smooth muscles, and PDE5 inhibitors treat erectile dysfunction ^45,46^. In addition, ENPP3 has activity in a broad pH range compared to ENPP1 which may thus provide ENPP3 an outsized advantage in the acidic tumor microenvironment of some tumor types. Future studies to understand the contexts in which ENPP1 and ENPP3 play roles as immune checkpoints as well as when one or the other should be targeted in cancer immunotherapy are therefore warranted.

## Materials and Methods

### Fractionation of *Enpp1^-/-^* mouse kidney

Kidney from *Enpp1^-/-^* mouse was isolated and flash-frozen in liquid nitrogen. On day of homogenization, kidney was thawed in 1 ml of hypotonic buffer (10 mM Tris pH 7.5, 150 mM sucrose including EDTA-free cOmplete protease inhibitor (Roche)) and homogenized using bead mill (5 cycles, 40 seconds on, 40 seconds off, Fisherbrand). Following homogenization, NaCl was supplemented to 150 mM and Tris to 50 mM pH 7.5. Digitonin was added to a final concentration of 0.25% to permeabilize the plasma membrane but leave the organelles intact. Following a 30-60 minute incubation with rotation at 4°C, all soluble proteins were thus separated by centrifugation at 80,000g for 1 hour at 4°C. The remaining pellet was solubilized using 1.5-2% NP-40 in 50 mM Tris pH 7.5 and 150 mM NaCl for 2-3 hours with rotation at 4°C. A second centrifugation at 80,000g for 1 hour at 4°C was performed to isolate the solubilized membrane proteins.

The supernatants from the centrifuge spins were diluted 50x into buffer containing 20 mM Tris pH 7.5 and 0.1% NP-40 prior to anion exchange on a 1-mL HiTrap Q column (Cytiva). The elution was performed from 0 mM to 500 mM NaCl in 20 mM Tris pH 7.5 with 0.1% NP-40 over a gradient involving 30 column volumes.

Following anion exchange, each fraction was evaluated for cGAMP degradation activity. 8µl of each fraction from AEX was supplemented with 1 µl of 10x physiological ion buffer, radioactive cGAMP and cold cGAMP to 5 μM final concentration for a total volume of 10 µl. At 1x concentration, physiological ion buffer contained 15 μM ZnCl_2_, 2.5 mM CaCl_2_, 1mM MgCl_2_, 100 nM MnCl_2_, 5 mM KCl, 150 mM NaCl, and 50 mM Tris pH 7.5. The reactions were incubated at room temperature overnight, after which the radioactive TLC described above was performed to evaluate the extent of cGAMP degradation in each fraction.

The fractions with the highest activity from AEX identified via TLC were pooled, concentrated and subjected to size exclusion on a Superose 6 Increase 10/300 GL column (Cytiva). Each fraction from size exclusion was again evaluated for activity. The entire fraction was subject to MS analysis at the Vincent Coates Foundation Mass Spectrometry laboratory, Stanford University Mass Spectrometry. 1555 proteins were identified in total (Supplementary Data Set). ENPP3 ranked #4 on the list.

### Synthesis and purification of ^32^P-GAMP

Radiolabeled ^32^P-ATP (3000 Ci/mmol) used for synthesis of ^32^P-cGAMP was purchased from Perkin Elmer. 1 μM purified sscGAS was incubated with 20 mM Tris-HCl pH 7.4, 2 mM ATP, 2 mM GTP, 20 mM MgCl2, and 100 μg/mL herring testis DNA (Sigma) for 1-3 days. Resulting ^32^P-cGAMP was purified via preparatory TLC as previously described.^1^

### 32P-cGAMP degradation TLC assay

To test for the presence of cGAMP hydrolases, 1 nM ^32^P-cGAMP and 5 μM cGAMP were incubated in fractions from anion exchange and size exclusion for 1-20 hours supplemented with physiological levels of ions (15 μM ZnCl_2_, 2.5 mM CaCl_2_, 1 mM MgCl_2_, 5 mM KCl, 100 nM MnCl_2_, 150 mM NaCl) with 50 mM Tris pH 7.5.

1-2 μL from the reaction was spotted on a silica TLC plate (Millipore Sigma) and allowed to dry for a minimum of 15 minutes. The TLC was developed in a solvent containing 85% ethanol and 5 mM NH_4_HCO_3._ Plates were dried and exposed to a Storage Phosphor Screen (Molecular Dynamics) overnight before being imaged with a Typhoon 9400 Imager (Molecular Dynamics).

### ATP degradation assay

ATP assays were conducted in 15 μL total volumes in a 384 well plate. 5 uM ATP were combined with organ lysate (0.1%) or recombinant ENPP3 (5 nM) and assay buffer (50 mM Tris pH 7.5, 150 mM NaCl, 500 μM CaCl2, 10 μM ZnCl2). Reactions were started at indicated times and ended simultaneously by heating at 95 °C for 10 minutes. 10 μL of each reaction were transferred to a white-walled 384 well plate, mixed with CellTiterGlo (5 μL), and luminescence was read after 15 minutes on a Tecan Spark plate reader.

### Cloning of full-length mENPP3

The DNA sequence encoding mouse ENPP3 (mENPP3) was cloned from a WT C57BL/6J mouse kidney. RNA from the mouse kidney was a generous gift from Wei Wei in the laboratory of Prof. Jon Long. A cDNA library was generated from the RNA using the iScript cDNA synthesis kit (BioRad). Full-length ENPP3 was PCR amplified using Phusion High-Fidelity DNA polymerase (Thermo). The PCR product was inserted into a pcDNA3 vector using Gibson assembly containing a TEV site followed by a TARGET-tag,^2–4^ 12 histidine residues and a stop codon as previously published.^4,5^

Primers for amplification of full length mENPP3 from cDNA:

Fwd: 5’-CTACGGGAACAATGGATTCCAG
Rev: 3’-CATTCAAATAATGGTTTCAAACGTGGGCAGATACGTC

Primers for amplification of full length mENPP3 for Gibson assembly:

Fwd: 5’-GGAGACCCAAGCTGGCTAGTTAAGCTTGCCATGGATTCCAGGCTAGCATTAGCCACAGAG
Rev: 5’–CTTACCTTGGAAGTACAGGTTCTCTCTAGAAATAATGGTTTCAAACGTGGGCAG

### Cloning of secreted mENPP3

cDNA from above was used to amplify all of mENPP3 except for its cytosolic and transmembrane domains (nucleotide 136-2622). The mENPP2 signal sequence was attached to the 5’ end of the mENPP3 sequence using successive rounds of PCR. The secreted mENPP3 construct was then inserted into a pcDNA3 vector using Gibson assembly containing a C-terminal TEV site followed by a TARGET-tag,^2–4^ 12 histidine residues and a stop codon as previously published.^4,5^

Primers for addition of mENPP2 signal sequence:

Fwd Primer 1: 5’-CTCTGCTTAGGAAGGAAACCTGAAGAACAAGGCAG
Fwd Primer 2: 5’-GGTAATATCCTTGTTCACTTTTGCCATCGGCGTCAATCTCTGCTTAGGAAGGAAACC
Fwd primer 3: 5’-GCCATGGCAAGACAAGGCTGTTTCGGGTCATACCAGGTAATATCCTTGTTCAC
Rev: 5’-AATAATGGTTTCAAACGTGGGCAGATACGTC

Primer for amplification of secreted mENPP3 for Gibson assembly:

Fwd: 5’ – TATAGGGAGACCCAAGCTGGCTAGTTAAGCTTGCCATGGCAAGACAAGGCT
Rev: 5’- CTTACCTTGGAAGTACAGGTTCTCTCTAGAAATAATGGTTTCAAACGTGGGCAGA

Cloning of mENPP3 mutations:

The H329A variant was introduced via site-directed mutagenesis. Mutated plasmids were cloned using pfuTurbo DNA polymerase (Agilent) and parent plasmids were degraded by DpnI (NEB). All plasmids were transformed into DH5α cells.

#### Primers for generation of H329A mutation

Fwd: 5’- CCTGATTCTGCAGGGGCGTCGAGTGGACCAGTC
Rev: 3’ – GACTGGTCCACTCGACGCCCCTGCAGAATCAGG

#### Primers for generation of T205A mutation

Fwd: 5’- GTATCCCACCAAAGCCTTCCCAAATCATTATAC
Rev: 3’ – GTATAATGATTTGGGAAGGCTTTGGTGGGATAC

### Cell culture

The HEK 293T cell line was procured from ATCC. The HEK 293T cGAS ENPP1 ^-/-^ cell line originates from previously published work.^4^ All cell lines were maintained in a 5% CO2 incubator at 37°C. 293T cell lines were maintained in DMEM (Corning Cellgro) supplemented with 10% FBS (Atlanta Biologics, v/v) and 100 U/mL penicillin-streptomycin (ThermoFisher).

### Quantification of cGAMP

293T cGAS ENPP1^-/-^ cells were plated in tissue culture treated plates coated with 2% PurCol (Advanced BioMatrix). 24 hours after transient transfection with mouse or human ENPP3 via Fugene 6 (Promega), the media was gently removed and replaced with serum-free DMEM supplemented with 1% insulin-transferrin-selenium-sodium pyruvate (ThermoFisher) and 100 U/mL penicillin-streptomycin. 24 hours after changing to serum-free media, media and cells were collected and centrifuged at 1000 rcf for 5 minutes at room temperature. Cells were washed with PBS. Cells were lysed in *M-PER* Mammalian Protein Extraction Reagent (Thermo Scientific) and analyzed for cGAMP content alongside media using the 2’3’ cGAMP ELISA kit (Cayman Chemical).

### Quantification of degradation of UDP-GlcNAc

0.8 and 4 nM of recombinant mENPP3 was incubated with 10 μM UDP-GlcNAc in buffer containing 50 mM Tris pH 7.5, 150 mM NaCl, 0.5 mM CaCl_2_, 10 uM ZnCl_2_, 0.1% NP-40 at room temperature in 20 μL reactions. The reaction was inactivated at 95°C at the indicated timepoints. Following the reaction, the UMP/CMP-Glo™ Glycosyltransferase Assay (Promega) was used to quantify the resulting UMP. Data was normalized to enzyme concentration used.

### Characterization kinetics of ENPP3

To characterize the biochemical kinetics of ENPP3, 7.5 nM of either mouse or human ENPP3 was incubated with the indicated concentrations of cGAMP spiked with ^32^P-cGAMP. Mouse ENPP3 was purified as described earlier; human ENPP3 was purchased from AcroBiosystems (#EN3-H52H4). Reactions were performed in buffer containing 20 mM Tris pH 9, 50 mM NaCl, 1 mM CaCl_2_, and 100 uM ZnCl_2_. Timepoints of 0, 10, 20, 30, 45, 60, 75, 90, 120 and 160 minutes were taken. At each timepoint, 1.5 μL from the reaction was spotted on a silica TLC plate (Millipore Sigma) and allowed to dry for a minimum of 15 minutes. The TLC was developed in a solvent containing 85% ethanol and 5 mM NH_4_HCO_3._ Plates were dried and exposed to a Storage Phosphor Screen (Molecular Dynamics) overnight before being imaged with a Typhoon 9400 Imager (Molecular Dynamics).

### Purification of secreted ENPP3 and ENPP1

Supernatant was supplemented with His-Pur cobalt resin (ThermoFisher), imidazole to 10 mM, NaCl to 150 mM, Tris to pH 7.5 and cOmplete protease inhibitor cocktail (Roche). Following incubation for 1 hour at 4°C, resin was washed with 15 column volumes of buffer made of 50 mM Tris pH 7.5, 150 mM NaCl (1x TBS) and 10 mM imidazole. A second wash of 15 column volumes of buffer made of 20 mM imidazole in 1x TBS was performed. ENPP3 or ENPP1 was eluted with 10 column volumes of 300 mM imidazole in 1x TBS. Buffer exchange to remove imidazole from the mixture was performed in centrifugal filter with a 10-kDa molecular mass cutoff (Millipore). Following quantification via micro-BCA assay (Thermo Fisher), ENPP3 or ENPP1 was stored in aliquots of 1 μM in 1x TBS with 0.1% NP-40 at -80°C.

### Purification of full-length ENPP3

Frozen cell pellets were resuspended in 10 mM Hepes pH 7.5, 150 mM sucrose and crushed in a bead mill (FisherScientific), 40 seconds on, 20 seconds off, for a total of 5 cycles. Following homogenization, the mixture was supplemented with Tris pH 7.5 to 50 mM, NaCl to 150 mM, imidazole to 10 mM, and cOmplete protease inhibitor cocktail (Roche). The mixture was centrifuged at 4°C at 20,000g for one hour and the supernatant was filtered through a .45 um syringe filter. The filtered supernatant was then purified and stored according to the protocol described for secreted ENPP3.

### Mouse Models

C57BL/6J (Stock #000664) and C57BL/6J-Sting1gt/J (Stock #017537) mice were purchased from the Jackson Laboratory. *Enpp3*^-/-^ and *Enpp3*^H329A^ mice were generated and characterized in house and bred with C57BL/6J-Sting1gt/J to generate *Enpp3*^H329A^ x *Sting*^-/-^ and *Enpp3*^-/-^ x *Sting*^-/-^ mice. Male and female mice were included in every experiment, unless otherwise noted. Mice were maintained at Stanford University in compliance with the Stanford University Institutional Animal Care and Use Committee regulations. All procedures were approved by the Stanford University Administrative Panel on Laboratory Animal Care.

Generation. First, single-guide RNAs (sgRNAs) were designed against the H329A locus in exon 11 of mouse Enpp3 using publicly available design tools (13). The sgRNA was then complexed with Alt-R S.p. Cas9 nuclease (Integrated DNA Technologies) as a ribonucleoprotein (RNP) particle. We then designed a donor sequence based on mouse Enpp3 to serve as the template for homologous recombination (Supplementary Table 3). The 200 nucleotide-long donor sequence contained blocking mutations near the PAM sequence to prevent repeated editing.

The donor sequence was then synthesized as single-stranded DNA (Integrated DNA Technologies). The RNP particles and donor template were microinjected into the pronuclei of one-cell embryos from C57BL/6 WT or ENPP1^H362A^ mice, which were then implanted into pseudo-pregnant mice. The injections were carried out by Stanford Transgenic, Knockout and Tumor Model Center. The initial litters of mice were backcrossed onto the WT C57BL/6 background and sequenced. Mice bearing the H329A mutation were crossed to each other to establish the ENPP3 H32A strain. Mice bearing a +1 frameshift mutation were crossed to each other to establish the ENPP3 KO strain.

Litter tracking. Male and female mice of indicated genotypes were set up in breeding pairs at 6-8 weeks of age. After an initial 2.5-week period, cages were checked twice a week for birth of new litters.

### Oligonucleotides for generating ENPP3^-/-^ and ENPP3^H329A^ mice

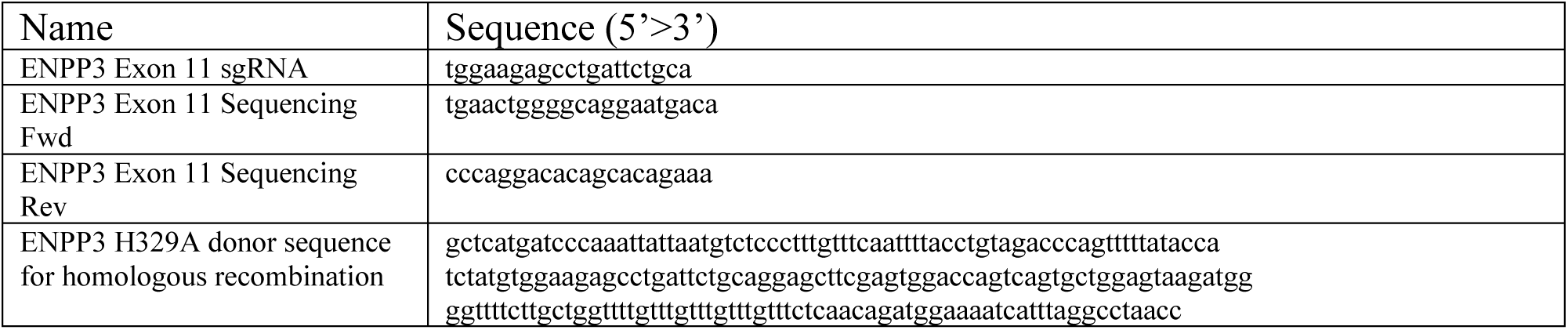

### Orthotopic E0771 tumor model

Five- to nine-week-old female mice were used for all experiments, and all tumor inoculations were performed with PBS as the vehicle. Mice of the indicated genotype were inoculated with 250,000 E0771 cells suspended in 100 μL into the fifth mammary fat pad. Tumor volumes were recorded and analyzed with a generalized estimation equation (width*height*height/2). Pairwise comparisons of the treatment groups at each time point were done by using post hoc tests with a Tukey adjustment for multiple comparisons. Animal death was plotted in a Kaplan–Meier curve with GraphPad Prism 9.3.1, and statistical significance was assessed by log-rank Mantel–Cox test.

### E0771 metastasis model

450,000 E0771.LMB.PuroR cells were per mouse injected into the indicated genotype via the tail vein using a 26-gauge needle. On day 30, lungs were harvested from mice, homogenized and plated in the presence of 1 μg/μL puromycin for 9 days to select for metastatic E0771 cells. Metastasis of each mouse was scored following staining of the E0771 cells by methylene blue and image analysis via Fuji Image J of blue-stained areas.

### B16 metastasis model

200,000 B16 cells were per mouse injected into the indicated genotype via the tail vein using a 26-gauge needle. On day 20, lungs were harvested from mice and fixed in formaldehyde. Surface metastases were counted and graphed using an unpaired t-test to calculate significance.

## Acknowledgements

We thank April Pawluk and the Arc Institute Scientific Publications Team for constructive feedback on the manuscript. We thank all Li Lab members for their constructive comments and discussion throughout the course of this study. R.M. was supported by National Science Foundation Graduate Research Fellowship Program Grant 1656518. S.W. was supported by the Stanford Medical Scholars Fellowship, Stanford Berg Scholars Program, Stanford Chi-Li Pao Foundation Alpha Omega Alpha Student Research Fellowship, and Arc Institute. G.C.A. is supported by Stanford University Medical Scientist Training Program grant T32-GM007365 and a Hertz Fellowship. This work was supported by NIH DP2CA228044 (L.L)., NIH 5R01CA258427-02 (L.L.)., and Arc Institute.

## Author Contributions

R.M. conceived the project, performed the experiments, analyzed results, coordinated the project, and wrote the research manuscript. S.W. performed the experiments, analyzed results, and wrote the research manuscript. J.A.C. and G.C.A. performed the experiments and analyzed results. X.L. performed the experiments. L.L. conceived the project, coordinated the project, wrote the research manuscript, and provided the project funding.

---

**Supplemental Figure 1:**
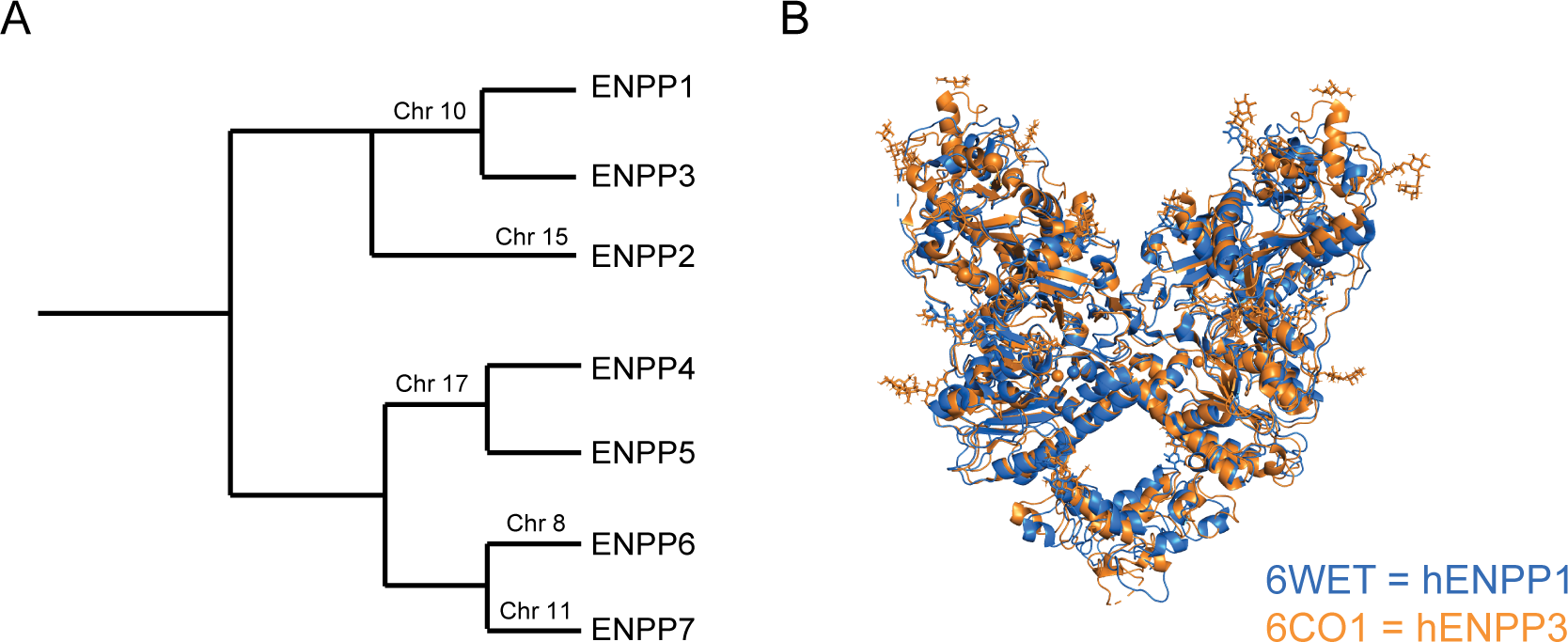
Sequence and structural homology between ENPP1 and ENPP3 (A) Evolutionary tree of the human ENPP family based on genetic similarity generated by Seaview. (B) Overlaid crystal structures of human ENPP1 (PDB 6WET) and human ENPP3 (PDB 6C01).

**Supplemental Figure 2:**
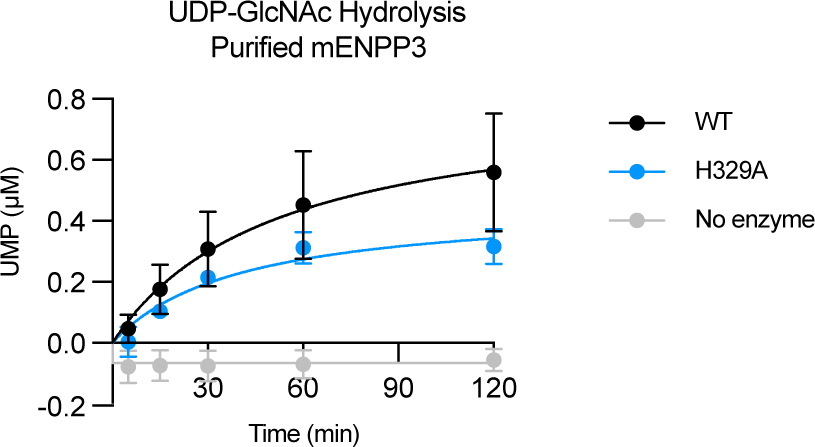
Kinetics of hydrolysis of UDP-GlcNAc by purified mouse WT and H329A ENPP3. Mean ± SEM (*n* = 4 for WT, *n* = 2 for H329A).

**Supplemental Figure 3:**
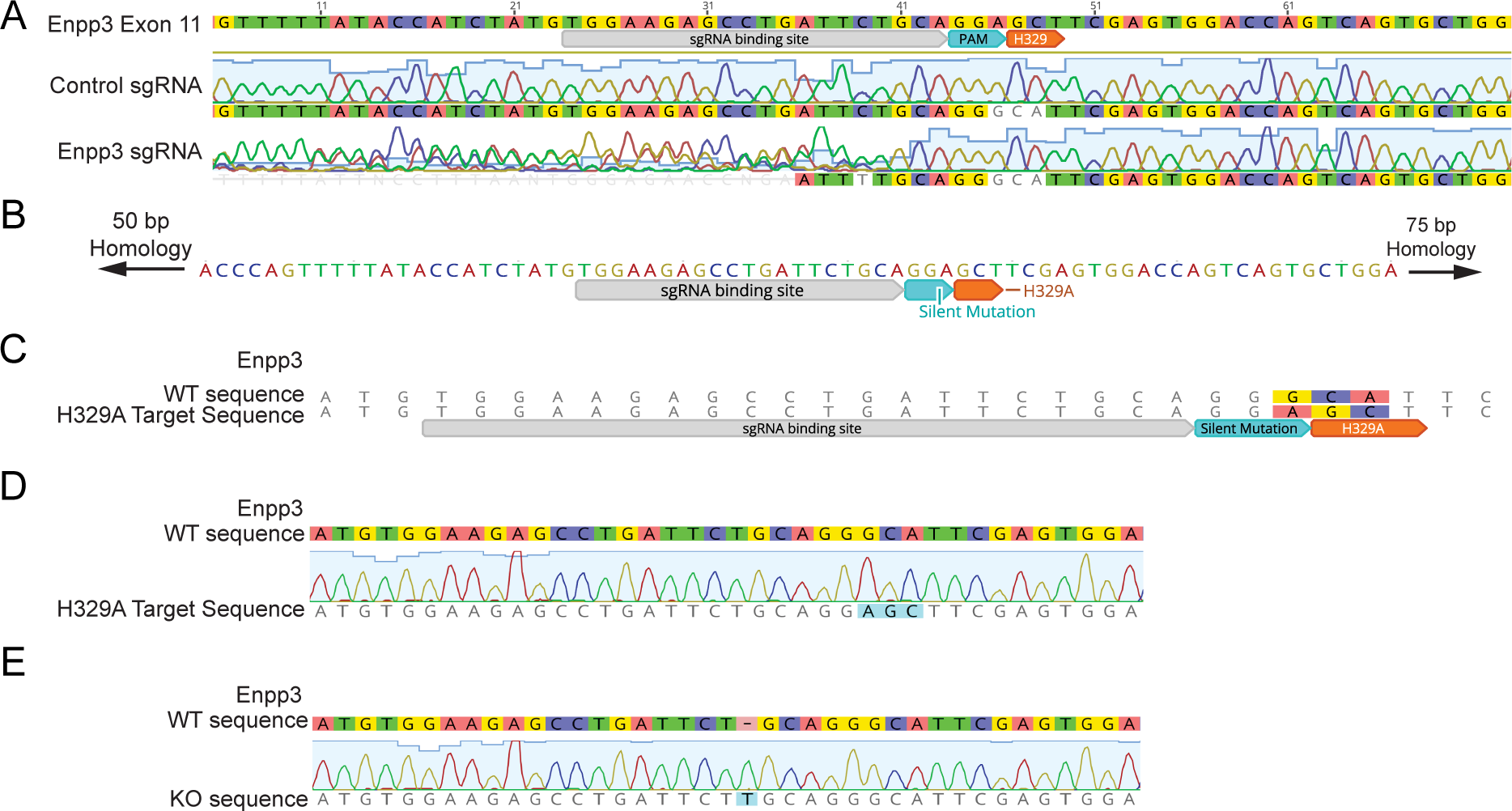
Generation of *Enpp3*^H329A^ and *Enpp3*^-/-^ mice. (A) A single guide RNA (sgRNA) was designed to target the region near H329A in exon 11 of *Enpp3*. The sgRNA and Cas9 were introduced into 4T1 cells through lentiviral transduction. The cells were then sequenced to determine editing efficiency at the intended cleavage site. (B) A homology arm was designed to generate the H329A point mutation through homologous recombination. A silent mutation was introduced downstream of H329 to prevent Cas9 from recognizing the PAM sequence following successful recombination. (C) Schematic illustrating the three base pair changes incorporated following successful recombination of the homology arm. (D) Schematic illustrating typical sequencing of genomic DNA from a homozygous *Enpp3*^H329A^ mouse. (E) Schematic illustrating typical sequencing of genomic DNA from a homozygous *Enpp3*^-/-^ mouse.

**Supplemental Figure 4:**
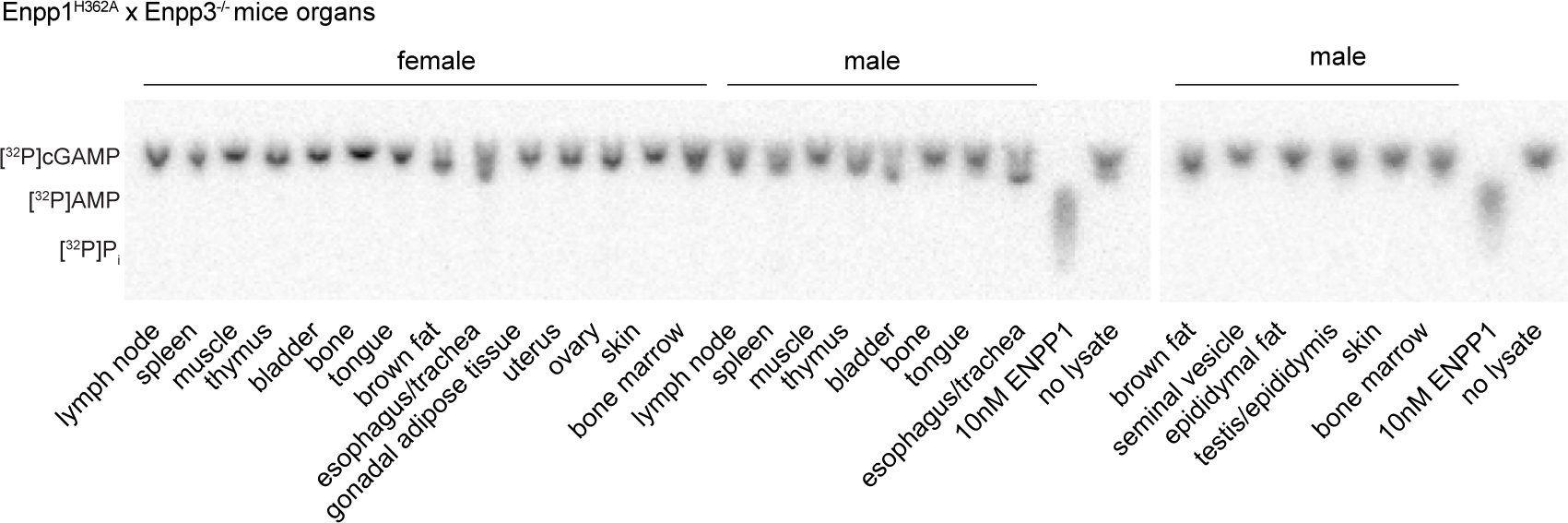
cGAMP hydrolase activity in indicated organ lysate of *Enpp1*^H362A^ x *Enpp3^-/-^* mice supplemented with physiological divalent ions (pH 7.5, 37°C, 24 hours).

**Supplemental Figure 5:**
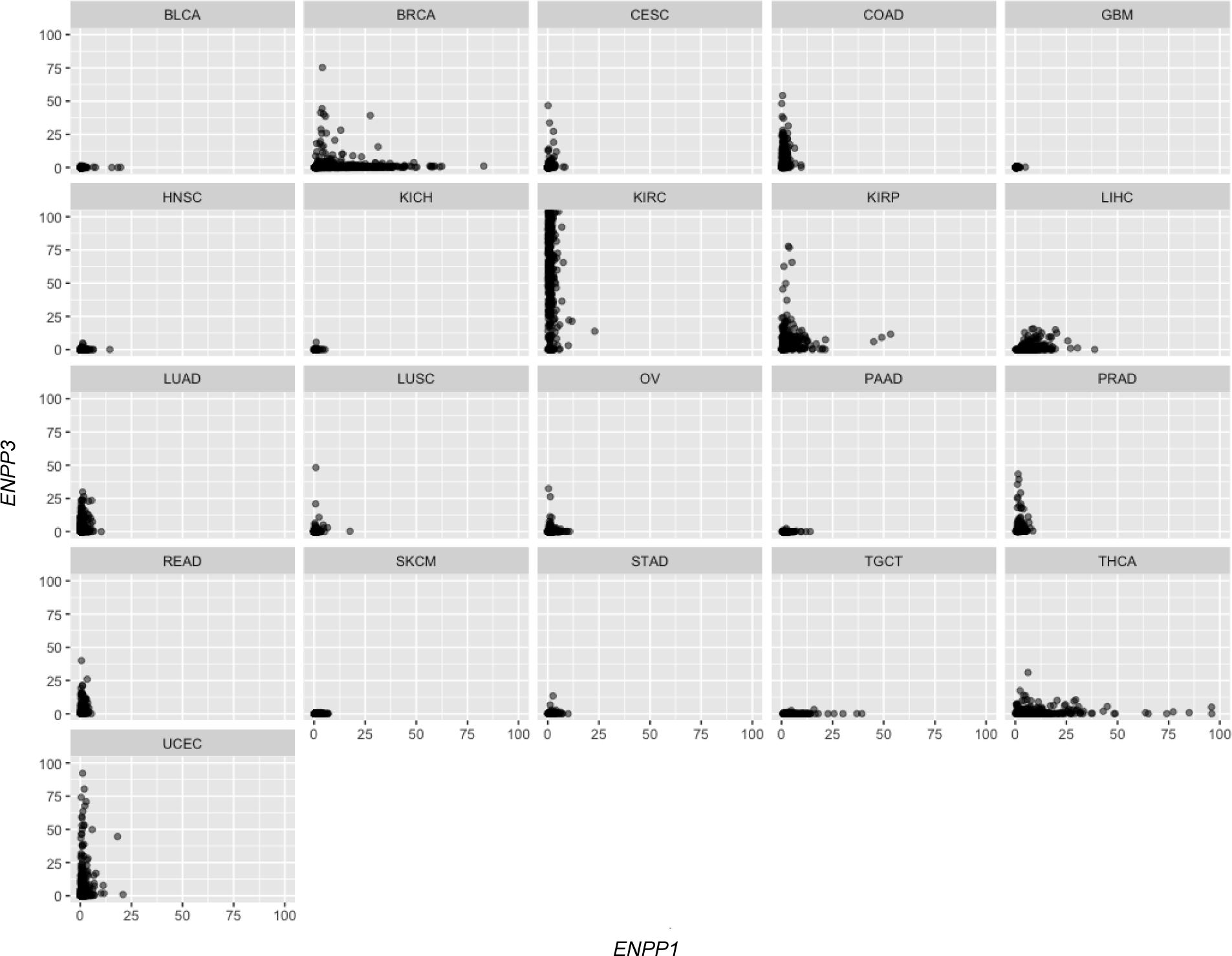
Quantification of expression of *ENPP1* and *ENPP3* from human cancer tissues in nTPM. Data from TCGA.

**Supplemental Table 1:** List of proteins and their abundances in fractionated *Enpp1*^-/-^ kidney lysate from proteomics mass spectrometry

